# Estimating abundance of a recovering transboundary brown bear population with capture-recapture models

**DOI:** 10.1101/2021.12.08.471719

**Authors:** Cécile Vanpé, Blaise Piédallu, Pierre-Yves Quenette, Jérôme Sentilles, Guillaume Queney, Santiago Palazón, Ivan Afonso Jordana, Ramón Jato, Miguel Mari Elósegui Irurtia, Jordi Solà de la Torre, Olivier Gimenez

## Abstract

Estimating the size of small populations of large mammals can be achieved via censuses, or complete counts, of recognizable individuals detected over a time period: minimum detected (population) size (MDS). However, as a population grows larger and its spatial distribution expands, the risk of under-estimating population size using MDS rapidly increases because the assumption of perfect detection of all individuals in the population is violated. The need to report uncertainty around population size estimates consequently becomes crucial. We explored these biases using the monitoring framework of the critically endangered Pyrenean brown bear that was close to extinction in the mid-1990s, with only five individuals remaining, but was subsequently bolstered by the introduction of 11 bears from Slovenia. Each year since 1996, the abundance of the population has been assessed using MDS and minimum retained (population) size (MRS), which corresponded to a reassessment of the MDS in the light of the new information collected in subsequent years (e.g., adding bears which were not detected the previous years but detected the current year). We used Pollock’s closed robust design (PCRD) capture-recapture models applied to the cross-border non-invasive sampling data from France, Spain and Andorra to provide the first published annual abundance and temporal trend estimates of the Pyrenean brown bear population since 2008. Annual population size increased fivefold between 2008 and 2020, going from 13 to 66 individuals. PCRD estimates were globally close to MRS counts and had reasonably narrow associated 95% Credibility Intervals. Even in cases where sampling effort is large compared to population size, the PCRD estimates of population size can diverge from the MDS counts. We report individual heterogeneity in detection that might stem from intraspecific home range size variation that result in individuals that move the most being most likely to be detected. We also found that cubs had a higher mortality rate than adults and subadults, because of infanticide by males, predation, maternal death, or abandonment. Overall, the PCRD capture-recapture modelling approach provides estimates of abundance and demographic rates of the Pyrenean brown bear population, together with associated uncertainty, while minimizing bias due to inter-individual heterogeneity in detection probabilities. We strongly encourage wildlife ecologists and managers to use robust approaches when researching large mammal populations. Such information is vital for informing management decision-making and assessing population conservation status.

## Introduction

Accurately and precisely estimating animal population size and trends over time is essential to inform conservation status and management decision-making (Nichols & Williams 2006). However, when animals, such as most large carnivores, are rare, elusive, solitary, largely nocturnal, highly mobile, and/or inhabiting large home ranges in remote and/or rugged habitats, population monitoring can be particularly challenging (Thompson 2013). Invasive physical tagging-based methods are difficult to implement, so population monitoring consequently often needs to rely on non-invasive sampling methods (Long & Zielinski 2008; Thompson 2013). Among them, molecular tools and camera trapping are now commonly used methods (e.g., Forsyth et al. 2022; Piel et al. 2022; Proctor et al. 2022). For species lacking unique natural individual patterns that can be identified from photos, non-invasive genotyping of DNA extracted from animal hair or scat often remains the most practical solution to estimate population abundance (Waits & Paetkau 2005).

Abundance of small populations of large mammals may be assessed using censuses or complete counts of unique individuals detected over a time period (Wilson & Delahay 2001; Keating et al. 2002), known as the minimum population size (Solberg et al. 2006; Miotto et al. 2007; Morin et al. 2022) and abbreviated here as MDS for minimum detected (population) size. In the case of genetic identification, MDS is defined as the number of unique genotypes identified among the genetic samples collected within the study area (Creel et al. 2003; Solberg et al. 2006). Obtaining a MDS via exhaustive counts, molecular sampling or extensive camera trapping, is often expensive, time consuming, and logistically demanding (Balme, Hunter & Slotow 2009; Blanc et al. 2013). In addition, as populations grow larger and spatial distributions expand, the risk of under-estimating population size using MDS increases sharply due to the rarely-fulfilled assumption of perfect detection of all individuals in the population (Solberg et al. 2006; Denes et al. 2015; Staton et al. 2022; Tourani 2022). The need to report uncertainty around population estimates consequently becomes crucial (e.g., Forney 2000; McGowan, Runge & Larson 2011). To address these issues, capture-recapture (CR) models are often used to estimate population abundance while accounting for the impossibility of detecting all individuals in a population (Otis et al. 1978). Whereas CR models were originally limited to live-trapping studies, they have been adapted for use with non-invasive DNA-based sampling (Lukacs 2005; Lukacs & Burnham 2005). In particular, non-invasive genetic CR models were specifically designed to account for issues such as individual identification errors due to genotyping errors, uncertainty in the date of individual detection, and the possibility of collecting multiple samples from the same individual across space within a single sampling occasion (Lukacs & Burnham 2005; Petit & Valière 2006; Lampa et al. 2013).

In standard closed-population CR models (whether or not they have been adapted to non-invasive genetic sampling), the population is assumed to be closed to changes in abundance both geographically (no immigration nor emigration) and demographically (no births nor deaths). Additionally, all individuals are assumed to have identical detection probabilities regardless of their individual attributes (e.g., age, body mass, social status) and habitat features (home-range location and composition) (Otis et al. 1978). However, these conditions are rarely fulfilled in real populations of wild mammals (e.g., Bellemain et al. 2005; Solbert et al. 2006).

In recent decades, considerable advances to these standard models have been developed to help alleviate issues linked to closure violation and detection heterogeneity (Lukacs & Burnham 2005). In particular, Pollock’s closed robust design (PCRD) CR modelling (Pollock 1982; Kendall, Nichols & Hines 1997) was developed in a maximum-likelihood (ML) framework to estimate survival, temporary emigration, and animal abundance while minimizing bias due to heterogeneity in detection among individuals. PCRD CR models rely on several so-called primary sampling occasions, each being composed of secondary occasions. The time interval between secondary sampling occasions must be short enough to meet the population closure assumption, while consecutive primary occasions should be sufficiently separated in time to allow the population to change. This framework is particularly suitable when population monitoring consists in periods of intensive sampling followed by a time gap with no or almost no sampling, like in the case of brown bear, *Ursus arctos* (almost no sampling during hibernation period in winter).

In the mid-1990s after decades of persecution, the brown bear population in the Pyrenees Mountains at the border of France, Spain and Andorra (Fig. 1) was on the edge of extinction with only five individuals remaining (Taberlet et al. 1997). Since then, the successful translocation of 11 bears from Slovenia (Quenette et al. 2019) has enabled the population to slowly recover. However, the fate of this critically endangered population (UICN France et al. 2017), isolated from the nearest Cantabrian brown bear population in north-western Spain by approximately 300 km, is still uncertain (Le Maho et al. 2013). Indeed, the population still displays a low abundance (MDS = 70 individuals in 2021; Sentilles et al. 2022), a low genetic diversity and a high consanguinity rate (expected heterozygosity He = 0.57 and consanguinity coefficient F = 0.13 in 2020; Bassi 2021), which make it particularly vulnerable to demographic, environmental and genetic stochasticity (Gilpin & Soulé 1986). Thus, developing reliable methods to accurately estimate population abundance and its trend over time is crucial to monitor the conservation status of this brown bear population and implement successful management plans.

**Figure 1:**
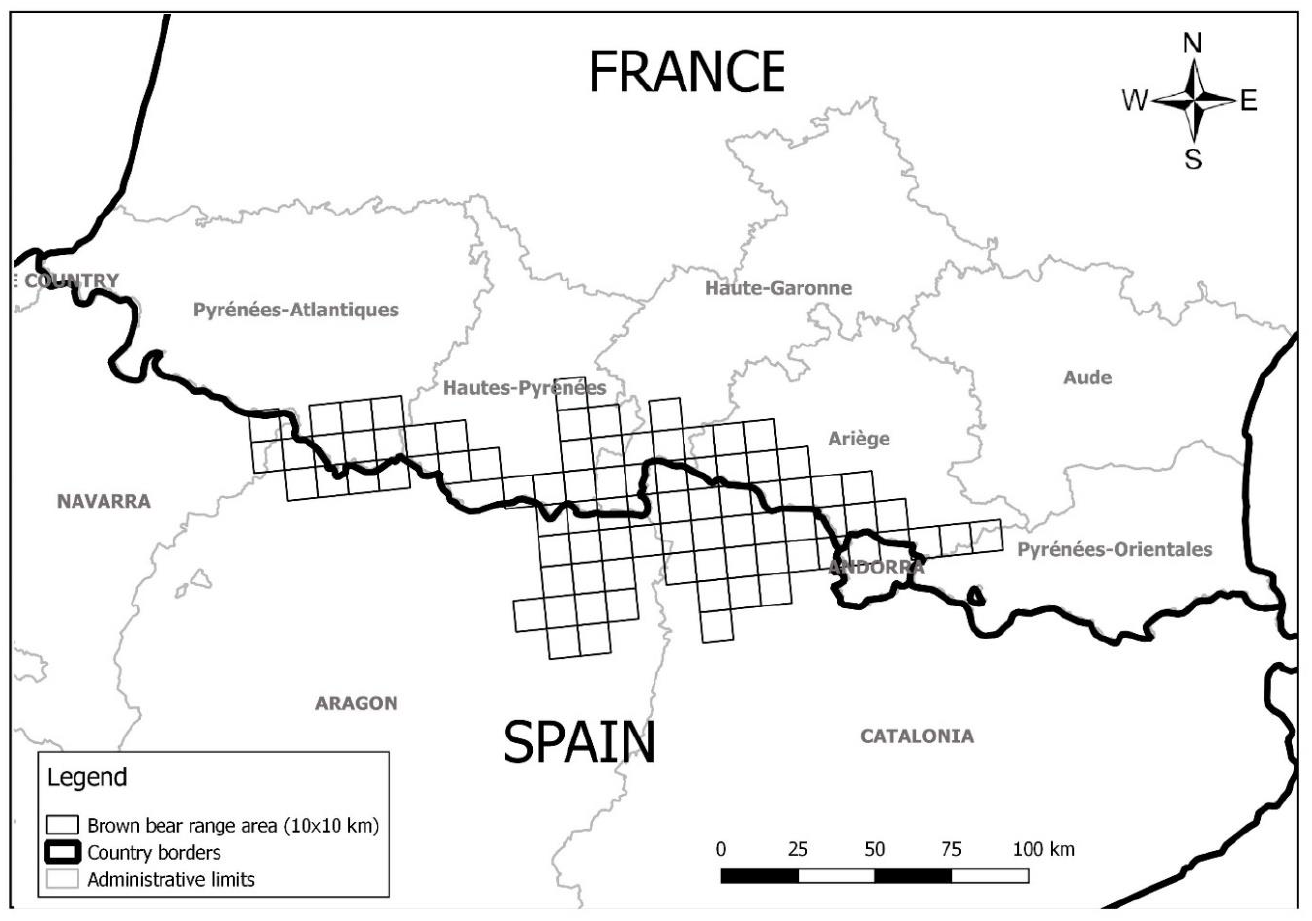
Map of the transboundary range area (on squares of 10 × 10 km) of the Pyrenean brown bear population for the year 2020.

Currently, non-invasive monitoring of the Pyrenean brown bear population relies on both systematic and opportunistic collections of bear presence signs (e.g., scats, hair, tracks, photos/videos, visual observations, attacks on livestock) in the Pyrenees Mountains combined with genetic or visual individual identifications (Sentilles, Vanpé & Quenette 2021; Sentilles et al. 2022). Similar to many large carnivore populations in Europe (Bischof, Brøseth & Gimenez 2016), the Pyrenean brown bear population is transboundary and occupies a politically and administratively fragmented landscape. It ranges indeed across the Principality of Andorra, two administrative regions (Occitanie and Nouvelle Aquitaine) divided across six counties in France, and three autonomous regions (Catalonia, Aragon and Navarra) and one Catalonian county with specific autonomous status (Val d’Aran) in Spain (Fig. 1). As such, cross-border multi-scale population monitoring cooperation (from national to local scales) is implemented to avoid overestimation, as individuals with home ranges overlapping borders may be detected in several political jurisdictions (Bischof, Brøseth & Gimenez 2016; Gervasi et al. 2016).

To date, the size of the Pyrenean brown bear population was annually assessed using MDS (Sentilles, Vanpé & Quenette 2021; Sentilles et al. 2022). However, this method assumes that all individuals present in the population have a detection probability of one. Because the population size was very small compared to the intensive sampling effort (Tables S1 and S2), the number of undetected individuals was assumed to be small. As the population was assumed to be geographically closed, the MDS of the current year was used each year to correct the MDS of previous years (e.g., to add bears which were not detected the previous years but detected the current year) and defined what we called the minimum retained (population) size, or MRS (Sentilles, Vanpé & Quenette 2021; Sentilles et al. 2022). MRS thus corresponded to a reassessment of the MDS in the light of the information collected in subsequent years. But although MRS could be regarded so far as a precise and accurate estimate of the true annual brown bear population size in the Pyrenees, it does not allow uncertainty assessment and MRS for year n is only available in year n+1 and sometimes needs a reassessment on year n+2 or n+3 (Sentilles et al. 2022). In adition, with increasing Pyrenean brown bear population size and range area, the number of undetected individuals over a year has increased over time. Finally, the outputs of demographic analyses of the Pyrenean brown bear population are used to inform management decision-making and policies (e.g., regulation, translocation, compensation). In this context, the reporting of abundance estimates and trends can be particularly prone to political influence (Darimont et al. 2018) and stakeholder skepticism. Therefore, implementing sound population monitoring tools and robust statistical methods to convey the uncertainty around abundance estimates is crucial. According to Lukacs and Burnham (2005), DNA-based CR methods provide the most useful methods to estimate abundance from small populations of up to a few thousand individuals, as in the Pyrenean brown bear population.

The aim of this study was therefore to provide the first published estimates of annual abundance of the Pyrenean brown bear population, while minimizing bias due to heterogeneity in detection probabilities among individuals. We used cross-border non-invasive sampling data collected from 2008 to 2020 in France, Spain and Andorra, for which individual identification was possible through genetic analyses or visual evidence combined with PCRD CR modelling. The development of new methods to estimate population abundance is timely, since it gives the possibility to compare the estimates obtained with the PCRD CR modeling approach with those from census approaches (MRS and MDS counts).

## Methods

### Brown bear population monitoring and bear sign collection

We carried out this study in the Pyrenees Mountains in southwestern Europe where the cross-border population of brown bears is present in the major part of the mountain range in France, Spain and Andorra and ranges over > 10,000 km^2^ in 2020 (Sentilles et al. 2022; Fig. 1). We used four different non-invasive methods to monitor the brown bear population in the French Pyrenees over the study period from 2008 to 2020 (Table S1):

1. Systematic trail walking (ST), equivalent to transect surveys (from 8 to 10 km long), spread homogeneously over the area of known, regular bear presence, which covers about 3,000 km^2^ in France (Vanpé et al. 2021; Sentilles et al. 2022; see Fig. S1). These transects were surveyed ten times (at least once per month) between May and November each year and bear signs were searched by two member teams of the Brown Bear Network (i.e. > 400 professionals and volunteers trained and managed by the bear team of the French Biodiversity Agency (OFB), thato is in charge of brown bear monitoring in the French Pyrenees by the French Minister of Ecology; Sentilles, Vanpé & Quenette 2021; Sentilles et al. 2022). To optimize bear detection, we set transects in the most favourable bear areas in terms of habitat quality and in bear transit areas detected using VHF and GPS collars or bear signs of presence. Transect staff searched for bear hair and scats on trails and in their immediate surroundings (see De Barba et al. 2010 for a similar approach). To improve the chances of finding hair samples, between five and seven hair traps were scattered along each trail. Each hair trap consisted of three small barbed wires fixed at three different heights onto a tree and where an attractive product (i.e. turpentine until 2016, beechwood tar called “smola” since 2017) was applied to encourage bear rubbing behavior (Berezowska-Cnota et al. 2017). Some of these hair traps were associated with a facing camera trap (similar to the camera trap methods described below) to help detect females with cubs and assess age class and number of individuals that rubbed on each focal tree, as well as the date of hair deposition (Parres et al. 2020).
2. Systematic by baited hair traps (SBHT) (2008 to 2011), corresponding to enclosures of about 20-30 m^2^ delimited by a strand of barbed wire fixed at a height of 50 cm (Woods et al. 1999; Kendall & McKelvey 2008; Quinn et al. 2022) and stretched around several trees. Bait consisting of a ~ 1-L mixture of rotten blood and fish was poured into the center of the area, with a reward of corn grains to increase recapture probability (see Woods et al. 1999; Castro-Arrellano et al. 2008; Gervasi et al. 2010). The trapping grid was established following designs and guidelines outlined in previous DNA-based inventories in North America (Mowat & Strobeck 2000; Boulanger et al. 2002) and average adult female home ranges of brown bears in the Pyrenees. The average home range size (Kernel 85%) of brown bears in the Pyrenees (excluding recently translocated individuals) was 84 km^2^ for adult females (N = 6) and 1,551 km^2^ for adult males (N = 6) (Halotel 2022; similar to the average home range of radio-collared adult bears in similar Eurasian regions: Huber & Roth 1993; Mertzanis et al. 2005; Gavrilov et al. 2015).We used a 4 × 4 km grid cell size based on known female range areas and a 8 × 8 km grid cell size for the remaining part of the study area, with one baited station placed in each grid cell. Hair traps were placed in the best predicted bear habitat, considering topography and accessibility by 4-wheel drive vehicles, a maximum of 10 min walk from the vehicle and bear expert opinion (tree types or tree species, with characteristics that make them more conspicuous for rubbing; González-Bernardo et al. 2021; Proctor et al. 2022). Sites were visited once every 15 days from May to September for sample collection and lure replacement.
3. Systematic by camera traps (SCT), corresponding to cameras (Leaf river Outdoor, HCO Soutguard SG 550 and Uway Nicht Trakker until 2013, and Bushnell Trophy Cam or NatureView HD and Reconyx HC600 or XR6 after 2013) equipped with movement detection that were fixed on trees in areas with frequent animal passage away from the walked transects and that were associated nearby with hair traps similar to the ones used for the systematic by trails method (Burton et al. 2015; Parres et al. 2020; see Fig. S1). Frequent animal passages were defined here as animals’ trails from all large mammals, which are visible in the vegetation and on the ground and that are often used by bears, as well as bear passage areas detected using VHF and GPS collars or bear presence signs. Each camera trap - hair trap station was visited once per month from April to November each year to collect samples and maintain cameras (Sentilles, Vanpé & Quenette 2021; Sentilles et al. 2022). We followed the same layout as above for SBHT protocol and placed one camera trap - hair trap station per cell. When hair samples could non-ambiguously be associated with photographs or videos, we analysed pictures in an attempt to individually identify bears based on natural markings, ear tags, or collars in order to avoid genetic analyses and decrease sampling costs.
4. Opportunistic monitoring (OM), corresponding to the opportunistic collection (i.e. with no specific sampling design) throughout potential bear range of all bear signs of presence gathered by any mountain users (e.g., hikers, foresters, hunters, skiers, fishermen, shepherds), as well as all putative bear damages on livestock and beehives (De Barba et al. 2010). Potential bear range is defined here as the areas surrounding bear presence, allowing random locations (for bear absences) to fall where bears could have visited (15 km from the edge of presence), as defined in Martin et al. 2012, and it covers > 10,000 km^2^. Bear signs include hair, scats, tracks, claw marks on trees, feeding clues, visual observations, etc. Feeding clues are carcasses of wild or domestic preys, overturning of a large stone, and anthill and bee or wasp swarms burst open. Mountain users report their observations to the bear team of the OFB. Testimonies are examined and approved by an expert from OFB. A conclusion as to its validity as bear evidence, “confirmed,” “probable,” “doubtful,” or “false,” is given to each putative bear presence sign that could be verified, on the same day or a few days after its transmission, according to the elements necessary for their verification (Sentilles et al. 2022). Bear observations are validated only if an indirect bear clue (scats, hair, footprints) is found at the sighting site or if a photo or video is provided by the observer. To confirm that eating clues are from brown bears, we specifically look for evidence of associated bear clues close by (e.g., footprints, claw marks, hair, scats). If the elements are not sufficient to make a decision or if the observer could not be found for the statement of his/her testimony, the evidence is classified as “impossible expertise”. Only confirmed bear clues are included in our analyses. Since 2014, verification of testimonies and damage reports have been occasionally carried out with the help of a scat-detection dog trained to search for brown bear scats (Sentilles, Vanpé & Quenette 2021). Only hair and scat samples collected during the same period (from May to November) as the ST systematic monitoring were included.

While all the four protocols (ST, SBHT, SCT, OM) were used in France, brown bear monitoring consisted of only the ST and OM protocols in Catalonia (Spain) and Andorra, and only the OM protocol in Aragon and Navarra (Spain). But note that this should not affect bear detection and population abundance estimation, since the choice of the monitoring methods was not dictated by the country or administrative unit but rather by the regularity of bear presence in the area (ST was implemented only in areas of known, regular bear presence in France, Spain and Andorra, while OM was implemented everywhere within the potential brown bear presence area). Although few individuals (mostly translocated animals and problematic bears) were temporally equipped with either VHF and/or GPS collars or ear tags over the study period, we analysed only the non-invasive sampling data. For all the four protocols, we paid particular attention when evaluating the date when the signs were left by the bears and discarded any sign for which uncertainty in the date was too high to define which month the bear was present (see Supplementary Materials provided in Supplementary Information). This study complies with the standards, laws, and procedures concerning animal research ethics of the countries, in which it was performed.

### Individual identification of bear signs

We used all validated non-invasive brown bear sign collected in the Pyrenees from 2008 to 2020 (Table S2) for which individual identification was possible. Individual identification of bears was primarily based on genetic analyses of hair (stored dry in envelopes) and scats (stored in microtubes filled with 96% ethanol) non-invasively collected in the field, as well as visual evidence (colouration, scars, GPS collars, or VHF ear tag transmitters) obtained by remote cameras when available (Sentilles et al. 2021). This visual identification was performed by bear experts from OFB and was validated only if a consensus was achieved among the experts.

Due to financial constraints, only a subset of all collected hair and scat samples were genetically analysed to identify individuals each year (mean ± SD = 35.16 ± 12.29, min = 17.5 in 2015 and max = 59.5 in 2008; Table S2). Samples that were sent to the lab each year were carefully selected so that we optimised the detection of individuals (e.g., we favoured samples from cubs of the year or subadults, as well samples that were collected in the expansion front of the population) and the genotyping success (e.g., freshest scats, avoidance of hair coming from different individuals).

Genetic samples were analyzed at the Laboratoire d’Ecologie Alpine (LECA) joint research unit from 2008 to 2012 using a multiple-tubes Polymerase Chain Reaction (PCR) approach (consisting in repeating each DNA amplification independently for each locus several times; Taberlet et al. 1996, 1997) and from 2013 to 2016 using high-throughput microsatellite genotyping on the Illumina platform (De Barba et al. 2017). From 2017 to 2020, samples were analyzed in our laboratory at ANTAGENE Company using a new multiple-tubes PCR approach (see the methods and Table S3 provided in Supplementary Information). In all cases, a minimum of four repeats for each sample was carried out to avoid genotyping errors associated with low quantities of DNA (Miquel et al. 2006). A total of 13 microsatellites markers and one (for LECA) to three (for our laboratory) sex markers were targeted by the multiplex PCR in order to identify individuals and assign gender (De Barba et al. 2017; see the methods and Table S4 provided in Supplementary Information). Further information on genotyping error rate and probability of identity-by-descent can be found in De Barba et al. (2017), Beaumelle (2016), Bassi (2021) and Table S4.

### Population abundance estimation using capture-recapture models

The results from all sources of individual identification (genetic analyses and tracking of natural or artificial marks) of all bear signs for which the month when the bear left the sign was known were then aggregated to compile a monthly detection history for each bear in the population from January 2008 to December 2020 (see Supplementary Materials in Supplementary Information). Data are available in Gimenez (2022).

We used a PCRD CR model (Pollock 1982; Kendall, Pollock & Brownie 1995; Kendall et al. 1997; see also in Williams, Nichols & Conroy 2022) to estimate population abundance. This method has been applied on a number of bear populations (Stetz et al. 2010; Pederson et al. 2012; McCall et al. 2013; Tosoni et al. 2017). PCRD CR models use a hierarchical sampling strategy, including widely-spaced “primary occasions”, between which the population is considered as open (i.e. with births, deaths and temporary emigration), and repeated captures in a short timeframe (called “secondary occasions”) between which the population is assumed to be closed to population changes. Data from secondary samples within each primary period are analyzed using closed models to derive estimates of detection probability and population size. Apparent survival and temporary emigration are estimated using open models by collapsing data from the secondary periods. Here, temporary emigration refers to some individuals that might temporarily emigrate to areas where foraging conditions or breeding success are better, or that might be temporarily unavailable for capture because they are in dens (Henle & Gruber 2017).

The population was assumed geographically closed, i.e. no emigration or immigration could occur between this population and another one outside the Pyrenees. We used years from 2008 to 2020 as primary occasions of capture (N = 13) and months from May to September as secondary occasions (N = 5), that is 65 occasions of capture in total. We chose these secondary occasions because no births occur in this time interval. We excluded months from October to April because of low activity of bears during hibernation and high mortality risks of cubs of the year during their first months of life (bear cubs are born in the den during January-February).

PCRD CR models allow estimating population abundance, detection probability and apparent survival while accounting for temporary emigration (Pollock 1982; Kendall, Nichols & Hines 1997). We accounted for temporary emigration with two parameters. First we used the probability of an individual being a temporary emigrant, given it was alive and present in the study area in the previous primary sampling occasion. The other temporary emigration parameter is the probability of an individual being a temporary emigrant given it was a temporary emigrant in the previous sampling occasion. There is no temporary emigration when both parameters are 0, random temporary emigration when both parameters are set and estimated equal (and the probability of an individual being present in the study area is not dependent on whether or not it was present in the study area in the previous sampling period) and Markovian temporary emigration when both parameters are set and estimated distinct (and the probability of an individual being present in the study area is conditional on whether it was present in the study area before). Apparent survival rate is the probability of surviving and staying in the study area, and is the product of true survival and fidelity to the study area.

We combined a frequentist with a Bayesian approaches to implement PCRD CR models. We first used a frequentist approach fitting 24 different models in total to estimate detection, survival and emigration parameters (Murray & Sandercock 2020). We considered four detection structures (constant, time-dependent considering variation between and within primary occasions and heterogeneous using finite mixtures, in which individuals may belong to one class of animals with a some detection probability in some proportion π or to another class of animals with a different detection probability in proportion 1 – π), two survival structures (constant and age-dependent using three age classes: i.e. cubs < 2 year old, subadults = 2-3 years old and adults > 3 years old) and three emigration structures (constant, random and Markovian) (see Table 1). We used the Akaike Information Criterion corrected for small sample size (AICc) to perform model selection (Burnham & Anderson 2002). These analyses were performed with the ‘RMark’ package (Laake 2013) that allows calling the Mark program (White & Burnham 1999) from R software (RCoreTeam 2013). Because we ran into boundary estimation issues, we then used a Bayesian approach to estimate annual population abundance, relying on the best supported model from the frequentist approach. PCRD CR models were first formulated in a Bayesian framework by Schofield and Barker (2011) and later by Rankin and collaborators (2016). However, it is only in the few last years that a Bayesian implementation of PCRD models has been made possible without ecologists having to code their own potentially complex sampling algorithms (Rankin et al. 2016; Riecke et al. 2018). These analyses were performed using program Jags (Plummer 2003; and Riecke et al. 2018 for PCRD models in particular). The rationale in considering both frequentist and Bayesian frameworks was to use the advantages of each of them: the Frequentist framework allows model selection via AICc without prohibitive computation time, and the Bayesian framework allows for obtaining interpretable estimates, improving estimation when sample sizes are low, and having access to full posterior conditional probabilities of model parameters. Scripts and codes are available in Gimenez (2022).

**Table 1:** Model selection from the frequentist capture-recapture approach using Pollock’s robust design (PCRD) capture-recapture (CR) modelling approach.

| Model | Survival structure | Detection structure | Emigration structure | AICc | ΔAICc | Model | Survival structure | Detection structure | Emigration structure | AICc |
| --- | --- | --- | --- | --- | --- | --- | --- | --- | --- | --- |
| 1 | Age-dependent | Heterogeneous | Random | 1496.43 | 0.00 | 1 | Age-dependent | Heterogeneous | Random | 1496.43 |
| 2 | Age-dependent | Heterogeneous | Markovian | 1496.90 | 0.47 | 2 | Age-dependent | Heterogeneous | Markovian | 1496.90 |
| 3 | Constant | Heterogeneous | Random | 1503.48 | 7.04 | 3 | Constant | Heterogeneous | Random | 1503.48 |
| 4 | Constant | Heterogeneous | Markovian | 1503.76 | 7.33 | 4 | Constant | Heterogeneous | Markovian | 1503.76 |
| 5 | Age-dependent | Heterogeneous | No | 1520.68 | 24.25 | 5 | Age-dependent | Heterogeneous | No | 1520.68 |
| 6 | Constant | Heterogeneous | No | 1528.73 | 32.30 | 6 | Constant | Heterogeneous | No | 1528.73 |
| 7 | Age-dependent | Time-dependent<br>(within primary occasions) | Random | 1548.00 | 51.57 | 7 | Age-dependent | Time-dependent<br>(within primary occasions) | Random | 1548.00 |
| 8 | Age-dependent | Time-dependent<br>(between primary occasions) | Random | 1548.28 | 51.85 | 8 | Age-dependent | Time-dependent<br>(between primary occasions) | Random | 1548.28 |
| 9 | Age-dependent | Time-dependent<br>(between primary occasions) | Markovian | 1548.58 | 52.14 | 9 | Age-dependent | Time-dependent<br>(between primary occasions) | Markovian | 1548.58 |
| 10 | Age-dependent | Time-dependent<br>(within primary occasions) | Markovian | 1549.03 | 52.60 | 10 | Age-dependent | Time-dependent<br>(within primary occasions) | Markovian | 1549.03 |
| 11 | Constant | Time-dependent<br>(within primary occasions) | Random | 1555.00 | 58.56 | 11 | Constant | Time-dependent<br>(within primary occasions) | Random | 1555.00 |
| 12 | Constant | Time-dependent<br>(between primary occasions) | Random | 1555.31 | 58.87 | 12 | Constant | Time-dependent<br>(between primary occasions) | Random | 1555.31 |
| 13 | Constant | Time-dependent<br>(between primary occasions) | Markovian | 1555.41 | 58.98 | 13 | Constant | Time-dependent<br>(between primary occasions) | Markovian | 1555.41 |
| 14 | Constant | Time-dependent<br>(within primary occasions) | Markovian | 1555.85 | 59.42 | 14 | Constant | Time-dependent<br>(within primary occasions) | Markovian | 1555.85 |
| 15 | Age-dependent | Constant | Random | 1562.58 | 66.15 | 15 | Age-dependent | Constant | Random | 1562.58 |
| 16 | Age-dependent | Constant | Markovian | 1562.86 | 66.43 | 16 | Age-dependent | Constant | Markovian | 1562.86 |
| 17 | Constant | Constant | Random | 1569.64 | 73.21 | 17 | Constant | Constant | Random | 1569.64 |

We compared PCRD estimates of the annual Pyrenean brown bear population abundance with both MDS and MRS counts. Note that MRS for 2020 is provisional and will be reassessed in the future (see above).

## Results

### Individual identification

From 2008 to 2020, we had 10,019 validated brown bear signs (e.g., hair, scats, tracks, visual observations, damages, photos / videos) collected throughout the Pyrenees year-round (Table S2). Among the 2,524 hair and scat samples which were sent for genetic analyses in France over this period, 1,648 (65%) allowed individual identification (Table S2). From 2008 to 2020, 98 different individuals (44 females, 41 males and 13 individuals with undetermined sex) were identified in the Pyrenees from May to September. Those individuals have been detected from 1 to 61 different capture occasions (median = 5.5, mean ± SD = 10.25 ± 12.23) over the study period from 2008 to 2020 (which include 65 occasions of capture in total).

### Model selection

The two top ranked models best supported by the data (with ΔAICc < 2) among the 24 fitted models both included age-dependent survival, heterogeneous detection, and either random or Markovian emigration (Table 1). All other models had much higher AICc (ΔAICc > 6; Table 1). Survival estimates of cubs, subadults and adults were nearly identical for both top ranked models (mean ± SE = 84.4 ±3.8%, 95.4 ± 2.8% and 96.2 ± 1.5%, respectively, except that the SE of the Markovian model is 2.9% instead of 2.8% as for the random model for subadults; Table 2). 72% of individuals had a low detection probability of 42%, whereas 28% of individuals had a high detectable probability of 85% (Table 2). The probability of leaving the study area was <10% for both models, whereas the probability of remaining outside the study area was 22% (Table 2).

**Table 2:**
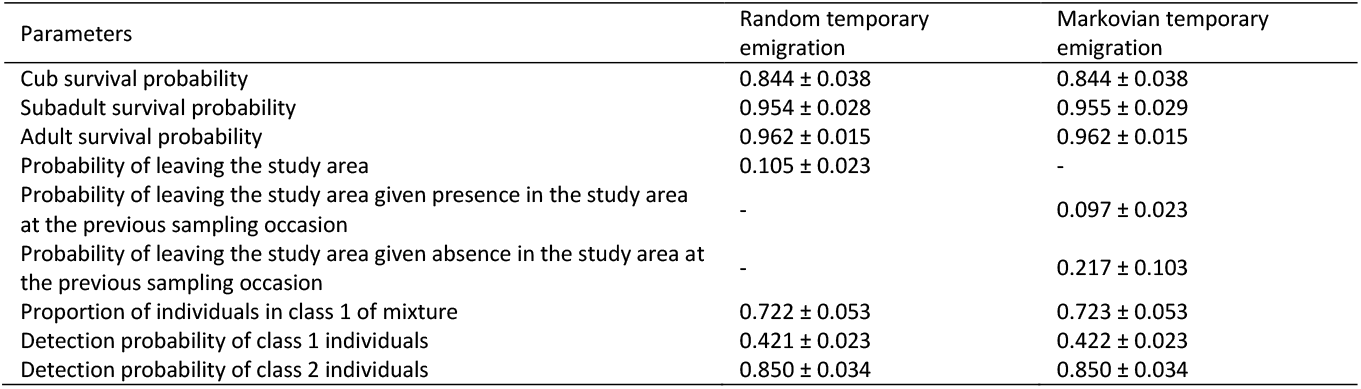
Parameter estimates (estimates ± SE) for the two best-supported models from the frequentist capture-recapture (CR) approach (see models 1 and 2 from Table 1) using a robust design, in which temporary emigration is either random (first column) or Markovian (second column).

| Parameters | Random temporary emigration | Markovian temporary emigration |
| --- | --- | --- |
| Cub survival probability | $0.844 \pm 0.038$ | $0.844 \pm 0.038$ |
| Subadult survival probability | $0.954 \pm 0.028$ | $0.955 \pm 0.029$ |
| Adult survival probability | $0.962 \pm 0.015$ | $0.962 \pm 0.015$ |
| Probability of leaving the study area | $0.105 \pm 0.023$ | - |
| Probability of leaving the study area given presence in the study area at the previous sampling occasion | - | $0.097 \pm 0.023$ |
| Probability of leaving the study area given absence in the study area at the previous sampling occasion | - | $0.217 \pm 0.103$ |
| Proportion of individuals in class 1 of mixture | $0.722 \pm 0.053$ | $0.723 \pm 0.053$ |
| Detection probability of class 1 individuals | $0.421 \pm 0.023$ | $0.422 \pm 0.023$ |
| Detection probability of class 2 individuals | $0.850 \pm 0.034$ | $0.850 \pm 0.034$ |

### Abundance estimation

Based on the best-supported model from the frequentist analysis (Table 2), we ran a Bayesian PCRD CR model, in which temporary emigration is random, survival is age-dependent survival and there is heterogeneity in the detection process. We used this model (see Table S5 for estimated parameters) to estimate annual abundance of the Pyrenean brown bear population. Bayesian PCRD estimates of the Pyrenean brown bear annual population abundance ranged from 13.0 with 95% credible interval (95% CI) = [12.8, 13.3] in 2008 to 66.2 with 95% CI = [64.8, 67.8] in 2020 (Fig. 2 and Table S6). We observed an increasing trend, with annual abundance displaying a fivefold rise between the beginning and the end of the study, with reasonably narrow 95% CI (Fig. 2 and Table S6).

**Figure 2:**
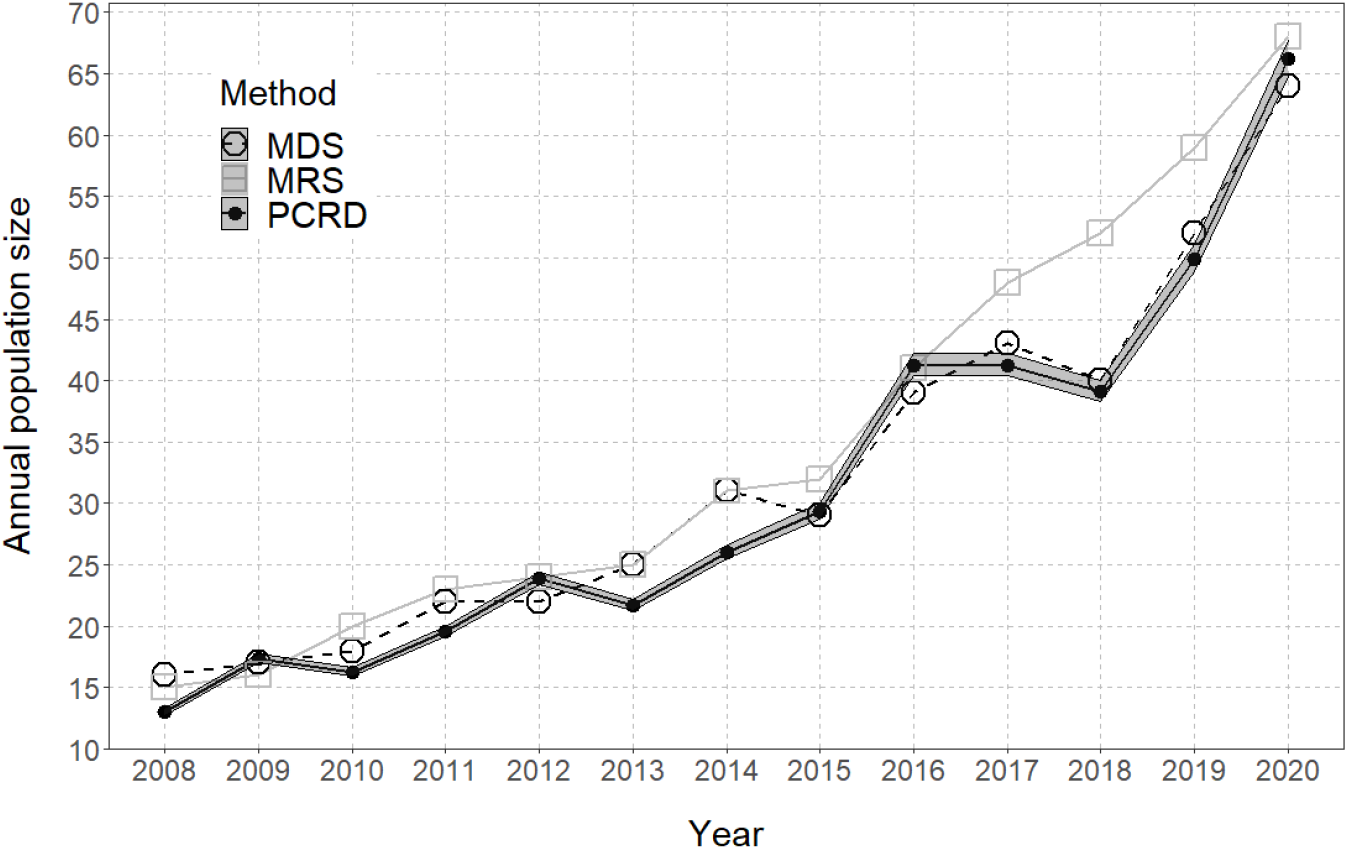
Variation in the annual population size of the Pyrenean brown bear from 2008 to 2020, estimated from Bayesian Pollock’s robust design capture-recapture approach (PCRD, black full circles and black full line, with the associated 95% credible interval in grey), compared to the Minimum Retained population Size (MRS, grey open squares and grey full line) and Minimum Detected population Size (MDS, black open circles and black dashed line) values. Note: MRS estimate for 2020 is provisional and probably slightly underestimated.

Differences in the estimates of the annual abundance of the Pyrenean brown bear population between the Bayesian PCRD CR modelling approach and census methods remained relatively small over the years (except from 2017 to 2019), with globally closer values between PCRD and MDS than between PCRD and MRS counts (mean difference ± SD = −1.02 ± 2.27 and −3.79 ± 3.95, respectively; Fig. 2 and Table S6). While PCRD estimates were either higher or smaller than MDS depending on the year, they were consistently smaller than MRS over the years except in 2009 and 2016 (+1.36 and +0.24, respectively; Fig. 2 and Table S6).

While MRS and MDS counts remained very close to each other before 2017 (mean difference ± SD = 0.89 ± 1.45), differences between MRS and MDS as well as between MRS and PCRD became much larger from 2017 (7.00 ± 3.56 and 7.65 ± 4.65, respectively; Fig. 2 and Table S6), considering that MRS for 2020 is provisional and will probably be reassessed upwards.

## Discussion

The Pyrenean brown bear population was shown to be composed of at least five individuals in 1995, indicating that population was then close to extinction (Taberlet et al. 1997). Topreserve the Pyrenean gene pool, increase genetic diversity and revive the population, the translocation of 11 bears from Slovenia was performed between 1996 and 2018 (Quenette et al. 2019). To assess the effectiveness of these conservation efforts and the current conservation status of the Pyrenean brown bear population, it is important to evaluate how population size has increased since the first translocations. We used PCRD CR models applied to the cross-border non-invasive sampling data from France, Spain and Andorra to provide the first published annual abundance estimates and trend of the critically endangered Pyrenean brown bear population from 2008 to 2020.

Our results suggest that the size of the Pyrenean brown bear population showed rapid population growth, displaying a fivefold rise between 2008 and 2020, reaching > 60 individuals (PCRD estimate = 66.2 with 95% CI = [64.8, 67.8]) in 2020. Most of the 11 translocations occured before 2008 (2 females in 1996, 1 male in 1997, 4 females and 1 male in 2006). Hence, the increase we observed in annual population size from 2008 to 2020 is not due to the introduction of new individuals into the population during the study period (1 male in 2016 and 2 females in 2018), but mainly to reproduction by an increasing number of individuals (Bassi 2021; Sentilles, Vanpé & Quenette 2021). The increase in the population abundance from 2018 cannot be explained by a female biased sex ratio, since the sex ratio among adults has been systematically biased towards females since 2012 (see Table S7). While this demographic success is encouraging for the short-term viability of the population, the fate of this critically endangered population is still uncertain due to high consanguinity, geographic isolation, fragmentation and small population size, which makes it particularly vulnerable to demographic, environmental and genetic stochasticity (Chapron et al. 2009; Le Maho et al. 2013; Beaumelle 2016; Bassi 2021).

We observed that PCRD estimates of the annual abundance of the Pyrenean brown bear population were close to MRS counts over all years except for 2017, 2018 and 2019, and had reasonably narrow 95% CI (Fig. 2 and Table S6). The fact that PCRD estimates are usually lower than MRS counts (and to a lesser extent, MDS counts) can be explained by the fact that our PCRD CR framework includes temporary emigration, which means that a bear that is not found during an entire year will not be included in the total population size estimate. Moreover, to use the PCRD CR framework, we excluded signs that were difficult to date, and those that fell outside of the secondary occasions (May to September), which left some individuals identified by MDS and MRS out of our database. Furthermore, MDS and MRS counts performed so far always included the individuals that were found dead in their yearly counts, while a PCRD CR model would only include them if the death occurred after the end of the primary occasion from October to December. Despite these limitations, our results suggest that the PCRD CR method provides reliable estimates of the size and trend of the Pyrenean brown bear population, while minimizing bias due to inter-individual heterogeneity in detection probabilities and quantifying sampling uncertainty surrounding these estimates.

The larger differences between MRS counts and both PCRD estimates and MDS counts in 2017 and 2018 (Fig. 2 and Table S6) may be partly explained by a limited number of DNA samples being collected in these years (N = 569 and 601, respectively) due to intensive translocation preparation efforts, compared for instance to 2015 and 2016 (N > 800; Table S2). This could result in a higher proportion of undetected individuals over the year, that could have been redetected during the following years. However, a large difference between MRS counts and both PCRD estimates and MDS counts was also observed in 2019 (9.08 and 7.00, respectively; Fig. 2 and Table S6), even though >800 DNA samples were collected over the year, among which 38% were analysed and 25% could be successfully genotyped (compared to 35.16 ± 12.29 % and 22.17 ± 7.22 %, respectively, on average from 2008 to 2020; Table S2). In addition, the difference between MRS counts and PCRD estimates was not positively correlated to the proportion of collected samples that were genetically analysed (F1,11 = 0.436, P = 0.52). The accentuation of the differences between MRS and MDS counts at the end of the study period (2020 excluded due to provisional MRS) thus likely indicates that we have now reached a point for which it becomes more and more difficult to detect all individuals within a year, even with intensive sampling and genotyping. As a consequence, the development of new metrics using capture-recapture methods to replace the MDS census approch to estimate the abundance of the Pyrenean brown bear populations is timely.

The model selection results highlighted two classes of individuals with significantly different detection probabilities (Table 2). A previous study on wolves highlighted the importance of accounting for individual heterogeneity in detection when estimating abundance of large carnivore populations (Cubaynes et al. 2010). Heterogeneity in the Pyrenean brown bears might stem from intraspecific home range disparities (McLoughlin, Ferguson & Messier 2000) making it more likely to find signs of individuals that move a lot, as well as from the fact that few bears were more easily visually identified due to their specific natural and/or artificial marks. The three individuals with long detection histories (N > 20 occasions) that were detected more frequently over the study period (> 85% of occasions) were indeed all large-sized adult males with particularly large home ranges and which were easily visually identified thanks to natural or artificial marks. Conversely, among the 10 individuals with long detection historyies (N > 20 occasions) that had the lowest detection probability (< 30% of occasions), we had both males and females and we did not observe any age effect. Natural and/or artificial marks (colouration, scars, GPS collars, or VHF ear tag transmitters) may have helped temporally or permanently identify some of the individuals of the population on photos or videos, causing potentially a bias in detection probabilities among individuals each month. However, this issue concerned only a few individuals each year (for natural marks: between 0 and 3 individuals according to years; for artificial marks: 2 individuals in 2008-2009, 0 in 2010-2015, 1 in 2016-2018, 4 in 2019 and 1 in 2020) and a few indices per individual (since natural marks are cryptic and not always visible on photos and videos). In the vast majority of cases, these individuals were detected independently each month via genetics on scats and hair. So we are confident this should not have significantly affected individual capture histories.

Another factor that might have caused heterogeneity in detection and might have affected the abundance estimate is the efficiency of people looking for bear sign. Some Pyrenean bears (e.g., dominant adult males and few adult females) displayed stable spatial behavior over the years (Camarra et al. 2015), making their movements predictable in time and allowing the researchers to become better at finding their signs (Fagen & Fagen 1996). Extending our approach to spatial capture-recapture (SCR) models that account for individual heterogeneity in the detection process by estimating individual-specific activity could help alleviate those issues (Royle et al. 2013; Borchers & Fewster 2016).

Interestingly, the PCRD CR modelling approach provides not only estimates of abundance but also estimates of demographic rates that cannot be provided by census approaches (MDS and MRS). We found an age-dependent effect on survival, with cubs surviving less well (84%) than subadults (95%) and adults (96%; Table 2). These results are consistent with previous estimates from Chapron et al. (2009) in the same population (0.77 ± 0.11 for cubs, 0.90 ± 0.09 for yearlings, 1.00 for sub-adults, and 0.97 ± 0.03 for adults in the Central sub-population between 1993 and 2005) and from cub survival estimates from most brown bear populations around the world (e.g., in British Columbia, Canada: 0.86 (0.74–0.96); McLellan 2015; in the Southern Scandinavian populations: 0.72; Swenson et al. 1997). In contrast, our cub survival estimate in the Pyrenees is much smaller than what was found in Northern Scandinavia (0.98; Swenson et al. 1997). However, cub mortality is known to vary widely among populations according to food availability, human disturbance and hunting management, with bear hunting either positively or negatively affecting cub survival depending on the population (Swenson et al. 2001). In the Pyrenees, bear hunting is prohibited and food availability is considered as good, but human disturbance can occur through various human activities including outdoor mountain activities, forestry, livestock farming, road traffic and hunting (Martin et al. 2012). The lower survival rate of cubs compared to other age classes was expected, since cubs are known to suffer from many mortality risks such as infanticide, predation, maternal death, or abandonment (Bunnell & Tait 1985) during their first year of life. In Scandinavia, about 80% of all cub mortality occurs during the mating season and is due to infanticide by males (Frank et al. 2017). While only a few infanticides, maternal deaths and abandonnementswere reported in the Pyrenees, their importance are probably greatly underestimated, since bear monitoring in the Pyrenees is mostly based on non-invasive methods. In addition, our estimate of cub survival is likely to be overestimated since our analyses do not take into account cub mortality at a very early age (< 4 months old) as we considered months from May to September as secondary occasions and births occur in the dens in January-February (Spady et al. 2007). As a result, some cubs may have died before we could even detect them for the first time. But cub mortality risks are not restricted to the first three months of their life and can also occur after April during late spring and summer.

In conclusion, our study shows that the PCRD CR modelling approach allows correcting for imperfect detection to provide estimates of abundance and demographic rates of the critically endangered Pyrenean brown bear population, while quantifying sampling uncertainty surrounding these estimates. Even in cases where sampling effort is large compared to population size, the PCRD CR abundance estimates can diverge from the minimum number known to be alive (MRS). In addition, MRS is obtained with at least one year’s delay, and the census approach is logistically and financially demanding. In the context of the demographic growth and geographical expansion of the Pyrenean brown bear population, we therefore recommend using our PCRD CR method rather than the former MDS metric to estimate the annual abundance and monitor the trend of this critically endangered population.

## Acknowledgements

We would like to acknowledge all field co-workers from the Brown Bear Network (ROB) and from Spanish and Andorran teams. We thank the editors and the three reviewers, Tim Coulson, Romain Pigeault and Susannah Woodruff, for their constructive comments and questions pertaining to the manuscript. We are very grateful to Tim Coulson and Marci Johnson for thorough English language editing and checking. We finaly acknowledge the recommender Nicolas Bech and the Managing editor of PCIEcology Marjolaine Hamelin. Preprint version 4 of this article has been peer-reviewed and recommended by Peer Community In Ecology (https://doi.org/10.24072/pci.ecology.100103).

## Data, scripts, code, and supplementary information availability

Data are available online: https://doi.org/10.5281/zenodo.7197316

Scripts and code are available online: https://doi.org/10.5281/zenodo.7197316

Supplementary information is available below as well as online: https://github.com/oliviergimenez/pyrenean-brown-bear-abundance/raw/master/Supplementary%20Information_Round2.pdf

## Supplementary Information

Supplementary Information for:

Estimating abundance of a recovering transboundary brown bear population with capture-recapture models

Cécile Vanpé, Blaise Piédallu, Pierre-Yves Quenette, Jérôme Sentilles, Guillaume Queney, Santiago Palazón, Ivan Afonso Jordana, Ramón Jato, Miguel Mari Elósegui Irurtia, Jordi Solà de la Torre, Olivier Gimenez

This part includes:

Supplementary Methods

Table S1 to S7

Figure S1

### Supplementary methods

#### Genetic analyses from 2017 to 2020

From 2017 to 2020, genetic analyses were conducted in our laboratory at ANTAGENE (https://www.antagene.com/en). DNA extraction was conducted according to a sterile process in a designated extraction room free of DNA. For each sample, disposable sterile tools were used and the bench was cleaned with bleach to avoid cross-contamination. Each sample was transferred to a sterile labelled microtube to proceed to DNA extraction. Sample tubes were surrounded by positive and negative extraction controls and lysed overnight at 56°C according to the manufacturer’s instructions (Nucleospin 96 Tissue Kit, Macherey-Nagel, Düren, Germany). DNA was isolated and purified using purification columns and vacuum filtration (Nucleospin 96 Tissue Kit, Macherey-Nagel, Düren, Germany). DNA was eluted with 100 μL of elution buffer to obtain final concentrations between 20-100 ng/μl. Extracts were stored in labelled 96-tube strip plates in a −20°C freezer.

For each DNA sample, 13 microsatellites and 3 sex identification markers (ZFX, 318.2 and SMCY) were amplified by two multiplex PCRs (polymerase chain reaction) four times and analyzed in two runs (one for each multiplex) with an automated sequencer (Table S6). Because the genetic sex marker described in the scientific publication De Barba et al. (2017) proved to be not very reproducible, the ANTAGENE laboratory uses a system of three pairs of primers allowing the amplification by PCR of two specific regions of the Y chromosome and one specific region of the X chromosome, according to a method developed and validated in all bear species (Bidon 2013). This system provides an internal positive control for all individuals, with the amplification of a region of the X chromosome present in males (XY) and in females (XX) and to amplify in duplicate a specific region of the Y chromosome present only in males (XY). This triple amplification guarantees an excellent recognition of the Y chromosome and therefore of males, and increases the reliability of characterization of the genetic sex, especially on DNA from degraded samples (hair, scats, etc.).PCR reactions were prepared step-by-step according to a unidirectional workflow starting in a clean room with positive air pressure to prepare sensitive reagents (enzymes and DNA primers) and continued in a pre-PCR room for combining DNA and reagents using filtered tips. Three negative and positive controls were included per PCR reaction. PCR amplifications were then performed in a dedicated post-PCR area in 96-well microplates at 10 μl final volumes containing 5 μl of mastermix Taq Polymerase (Type-It Microsatellite PCR Kit, Qiagen, Hilden, Germany), and either 0.80 μL of a first pool of 8 pairs of primers or 0.36 μl of a second pool of 8 pairs of primers at a concentration from 0.08 to 0.60 μM each, and a mean of 30 ng of genomic DNA (Table S6). Each pair of primers was coupled with a fluorescent dye (Table S6). Our PCR thermal protocol consisted of 95°C for 15 min, followed by 8 touchdown cycles of 95°C for 30 s, 62°C to 55°C for 90 s (decreasing 1°C per cycle), and 72°C for 30 s, then followed by 35 cycles of 95°C for 30 s, 55°C for 90 s, and 72°C for 30 s, ending with an extension of 60°C for 30 min. PCR products were resolved on an ABI PRISM 3130 XL capillary sequencer (ThermoFisher Scientific, Waltham, Massachusetts) under denaturing conditions (Hi-DiTM Formamide, ThermoFisher Scientific, Waltham, Massachusetts) with an internal size marker prepared once and dispatched equally in all sample wells of each multiplex run. The four electropherograms for each sample were analyzed using GENEMAPPER 4.1 (ThermoFisher Scientific, Waltham, Massachusetts) and analyzed independently by two analysts to determine the allele sizes for each marker of each individual. When the genotypes determined by each analyst did not agree, the electropherograms were read again, reading errors were resolved, and in case of persistent disagreement, ambiguous results were considered as missing data.

#### Dating of bear signs

For photos and videos, we used the metadata from the automatically triggered camera traps or cameras to define accurately the date of bear presence. For hair collected on baited hair traps, we used photo data collected on camera traps set up in front of baited hair traps when available to identify date when hair were left. From those specific bear signs, month of bear presence could be determined accurately based on the date when signs were left.

For other types of bear signs, we could not know precisely the date when signs were left and we relied on an evaluation of the time period when sign could have been left by the bear. More specifically, when hair collected on baited hair trap were not associated with any photo or video, we considered that the bear had left the hair during the time period included between the date of the last visit of the hair trap when barbed wire was cleaned and the date of the visit when hair were collected. If this time period was larger than 2 months, we discarded the hair sample from our analyses. We also discarded hair samples collected spontaneously outside systematic monitoring design, because the time interval during which they might have been left by the bear could not be evaluated precisely (bear hair deteriorates very slowly in the field), except in the case hair were associated with damage to livestock or beehives, in which case the estimated date of the damage provided the estimated date of hair deposition. Finally, we estimated the time interval when scats were dropped (≤2 weeks) by evaluating the freshness of the scat when collected in the field, using expert judgement in relation to the color and appearance of the scat, recent weather conditions (rain, sunshine, snow, temperature, etc.) and type of habitat (directly exposed to sun, under vegetation cover, etc.) (e.g., Sergiel et al. 2020 for a similar approach). When the time period during which hair or scat could have been left overlapped two different months, we considered as a proxy the month of the median date between maximum and minimum date of the time period as the month of bear presence, since this should not affect much our estimation of population size with capture-recapture analyses. Note that we selected preferentially fresher scats (with less DNA degradation) to send to the molecular laboratory, allowing a better genotyping success and identifying more individuals genetically (Sentilles, Vanpé & Quenette 2021). In France, we collected in total 4,022 hair or scat samples from 2008 to 2020, among which about 5.5% were excluded from our analyses due to inaccurate dating.

#### Compilation of monthly detection history of bears

Matching genotypes were considered to arise from the same individual and classified as recaptures as the combined non-exclusion probability of the 13 microsatellites for independent individuals and for sibships were negligeable (Lukacs & Burnham 2005). Importantly, we did not consider location data from GPS collar or VHF transmitters to compile detection history to avoid large inter-individual differences in monitoring pressure between bears, since it concerns respectively 5 bears and 1 bear for a period ranging from several months to a few years. Orphan cubs that were captured in the field and kept in captivity for a while for care before being released in the wild were considered as still present and detected in the population during the months of captivity (this concerns only 1 orphan cub during two months of captivity). For individuals for which we knew the date of death (N = 9), we used this information and right censored them in the corresponding detection histories. For translocated bears originating from Slovenia (N = 3), the first month of potential detection was the month of release in the Pyrenees.

---

**Table S1:**
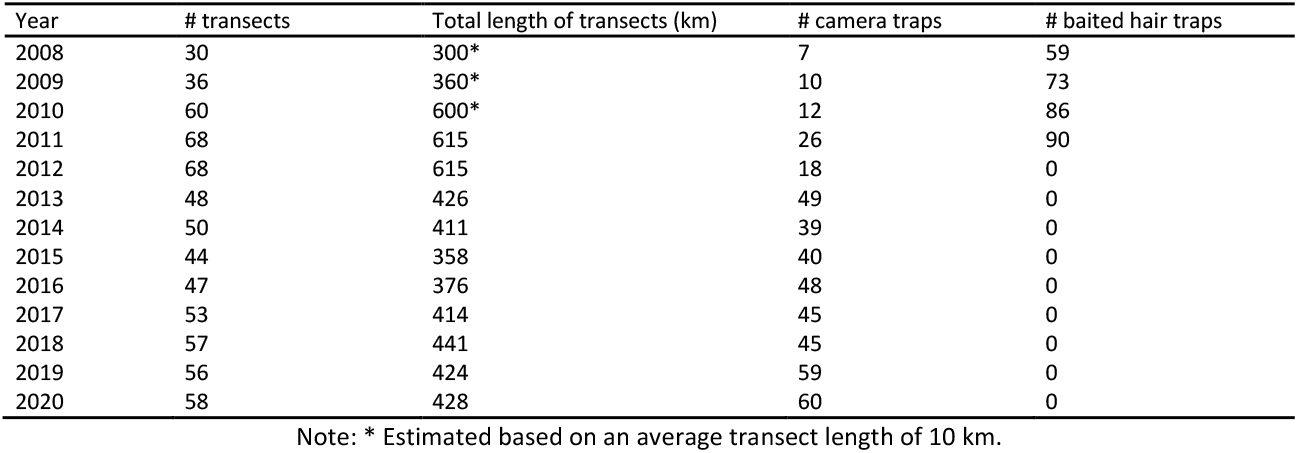
Systematic monitoring effort in the French Pyrenees in terms of number of transects (including 6 hair traps per transect in average), total length of transects (km), number of camera traps, number of baited hair traps per year between 2008 and 2020.

**Table S2:**
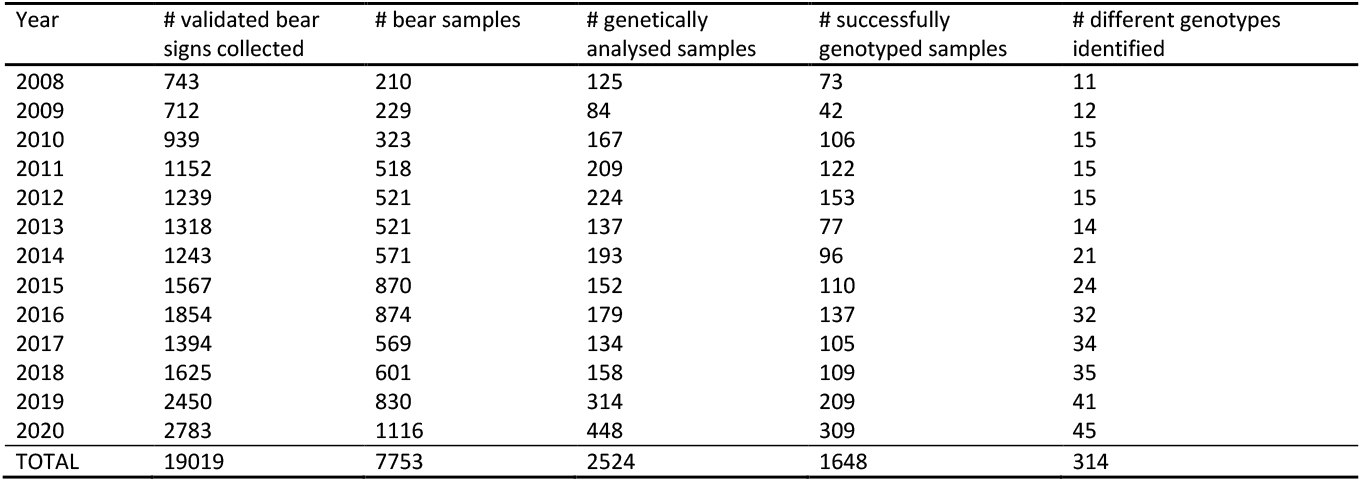
Total number of validated non-invasive brown bear signs (e.g., scats, hair, tracks, visual observations, damages, photos / videos) collected in the Pyrenees, total number of validated brown bear samples (i.e. scats and hair) collected in the Pyrenees, number of samples (among collected sampled) genetically analysed by the French molecular laboratory LECA or ANTAGENE, number of brown bear samples (among analysed samples) successfully genotyped and number of different brown bear genotypes identified (among successfully genotyped samples) per year between 2008 and 2020.

**Table S3:**
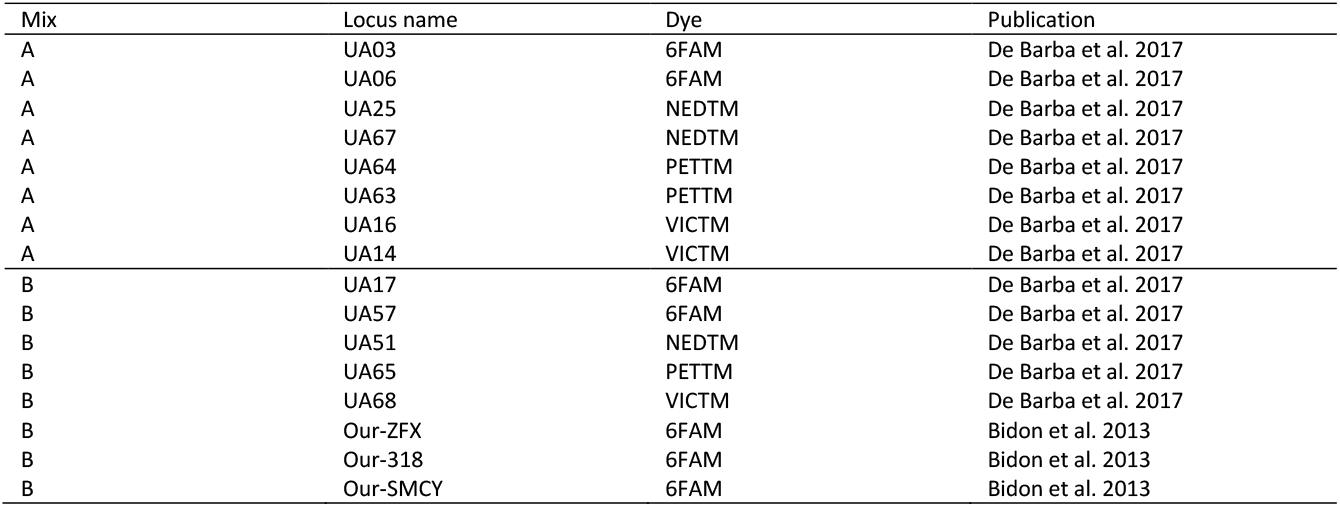
Combination of microsatellite markers used in each PCR mix and type of fluorescent dye used for each microsatellite marker from 2017 to 2020.

| Mix | Locus name | Dye | Publication |
| --- | --- | --- | --- |
| A | UA03 | 6FAM | De Barba et al. 2017 |
| A | UA06 | 6FAM | De Barba et al. 2017 |
| A | UA25 | NEDTM | De Barba et al. 2017 |
| A | UA67 | NEDTM | De Barba et al. 2017 |
| A | UA64 | PETTM | De Barba et al. 2017 |
| A | UA63 | PETTM | De Barba et al. 2017 |
| A | UA16 | VICTM | De Barba et al. 2017 |
| A | UA14 | VICTM | De Barba et al. 2017 |
| B | UA17 | 6FAM | De Barba et al. 2017 |
| B | UA57 | 6FAM | De Barba et al. 2017 |
| B | UA51 | NEDTM | De Barba et al. 2017 |
| B | UA65 | PETTM | De Barba et al. 2017 |
| B | UA68 | VICTM | De Barba et al. 2017 |
| B | Our-ZFX | 6FAM | Bidon et al. 2013 |
| B | Our-318 | 6FAM | Bidon et al. 2013 |
| B | Our-SMCY | 6FAM | Bidon et al. 2013 |

**Table S4:**
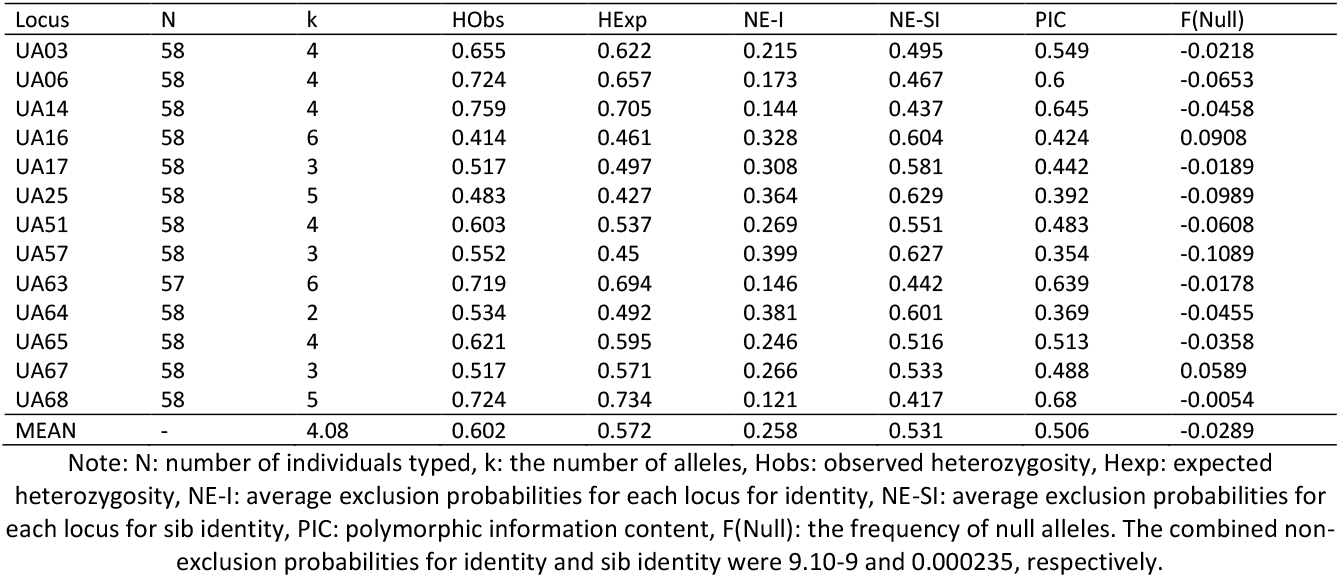
Summary statistics of the 58 different genotypes found in the Pyrenean brown bear population in 2020 for each of the 13 microsatellite loci provided by the allele frequency analysis of CERVUS software (Marshall et al. 1998).

**Table S5:**
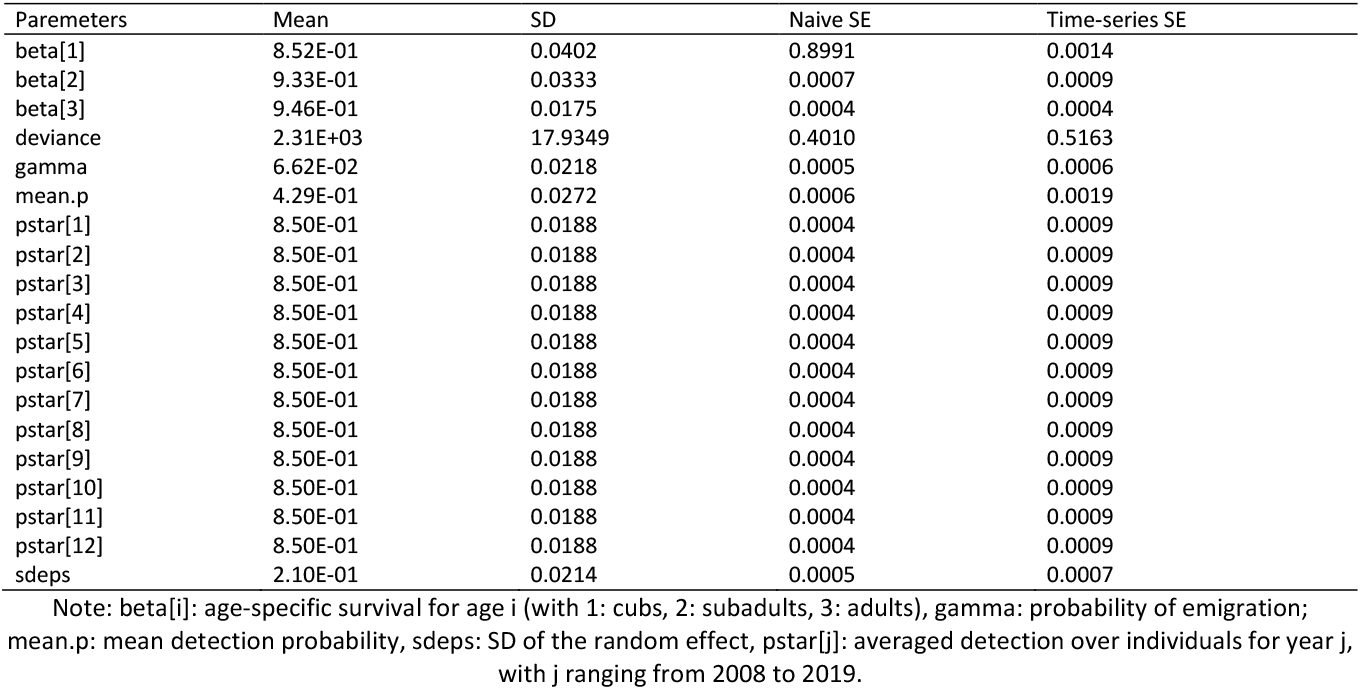
Parameters of the model in which temporary emigration is random, survival is age-dependent and there is heterogeneity in the detection process, estimated using a Bayesian robust-design capture-recapture (CR) approach.

**Table S6:**
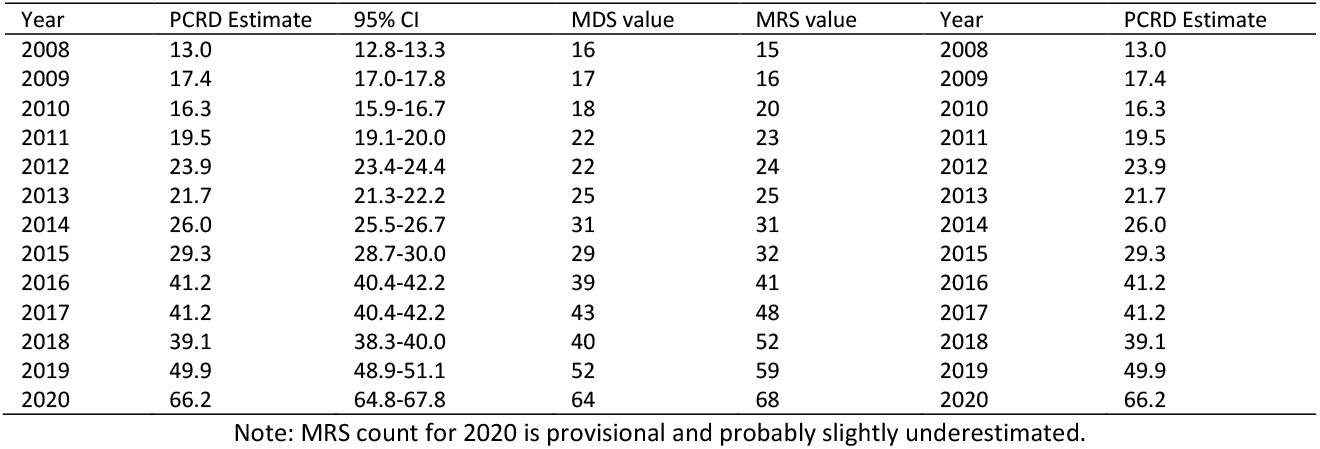
Comparison of the annual abundance of the Pyrenean brown bear population, estimated from Bayesian Pollock’s robust design (PCRD) capture-recapture (CR) approach (with associated 95% Credible Interval), with Minimum Detected Size (MDS, total number of different individuals detected in the population during the year) and Minimum Retained Size (MRS, reassessment of the MDS in the light of the information collected in subsequent years) values from 2008 to 2020.

| Year | PCRD Estimate | 95% CI | MDS value | MRS value | Year | PCRD Estimate |
| --- | --- | --- | --- | --- | --- | --- |
| 2008 | 13.0 | 12.8-13.3 | 16 | 15 | 2008 | 13.0 |
| 2009 | 17.4 | 17.0-17.8 | 17 | 16 | 2009 | 17.4 |
| 2010 | 16.3 | 15.9-16.7 | 18 | 20 | 2010 | 16.3 |
| 2011 | 19.5 | 19.1-20.0 | 22 | 23 | 2011 | 19.5 |
| 2012 | 23.9 | 23.4-24.4 | 22 | 24 | 2012 | 23.9 |
| 2013 | 21.7 | 21.3-22.2 | 25 | 25 | 2013 | 21.7 |
| 2014 | 26.0 | 25.5-26.7 | 31 | 31 | 2014 | 26.0 |
| 2015 | 29.3 | 28.7-30.0 | 29 | 32 | 2015 | 29.3 |
| 2016 | 41.2 | 40.4-42.2 | 39 | 41 | 2016 | 41.2 |
| 2017 | 41.2 | 40.4-42.2 | 43 | 48 | 2017 | 41.2 |
| 2018 | 39.1 | 38.3-40.0 | 40 | 52 | 2018 | 39.1 |
| 2019 | 49.9 | 48.9-51.1 | 52 | 59 | 2019 | 49.9 |
| 2020 | 66.2 | 64.8-67.8 | 64 | 68 | 2020 | 66.2 |

Note: MRS count for 2020 is provisional and probably slightly underestimated.

**Table S7:**
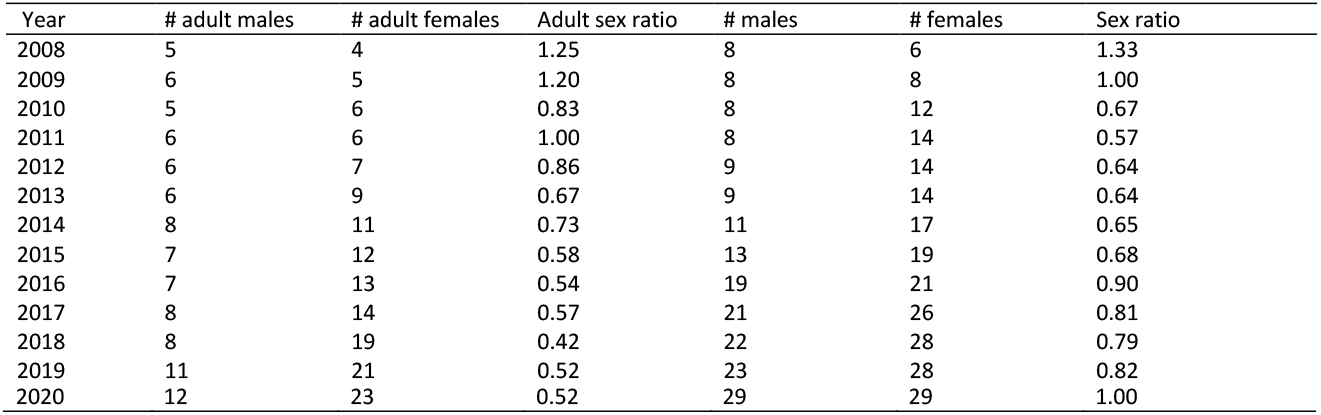
Evolution of the sex ratio of the Pyrenean brown bear population from 2008 to 2020 among all individuals and among adult only.

**Figure S1:**
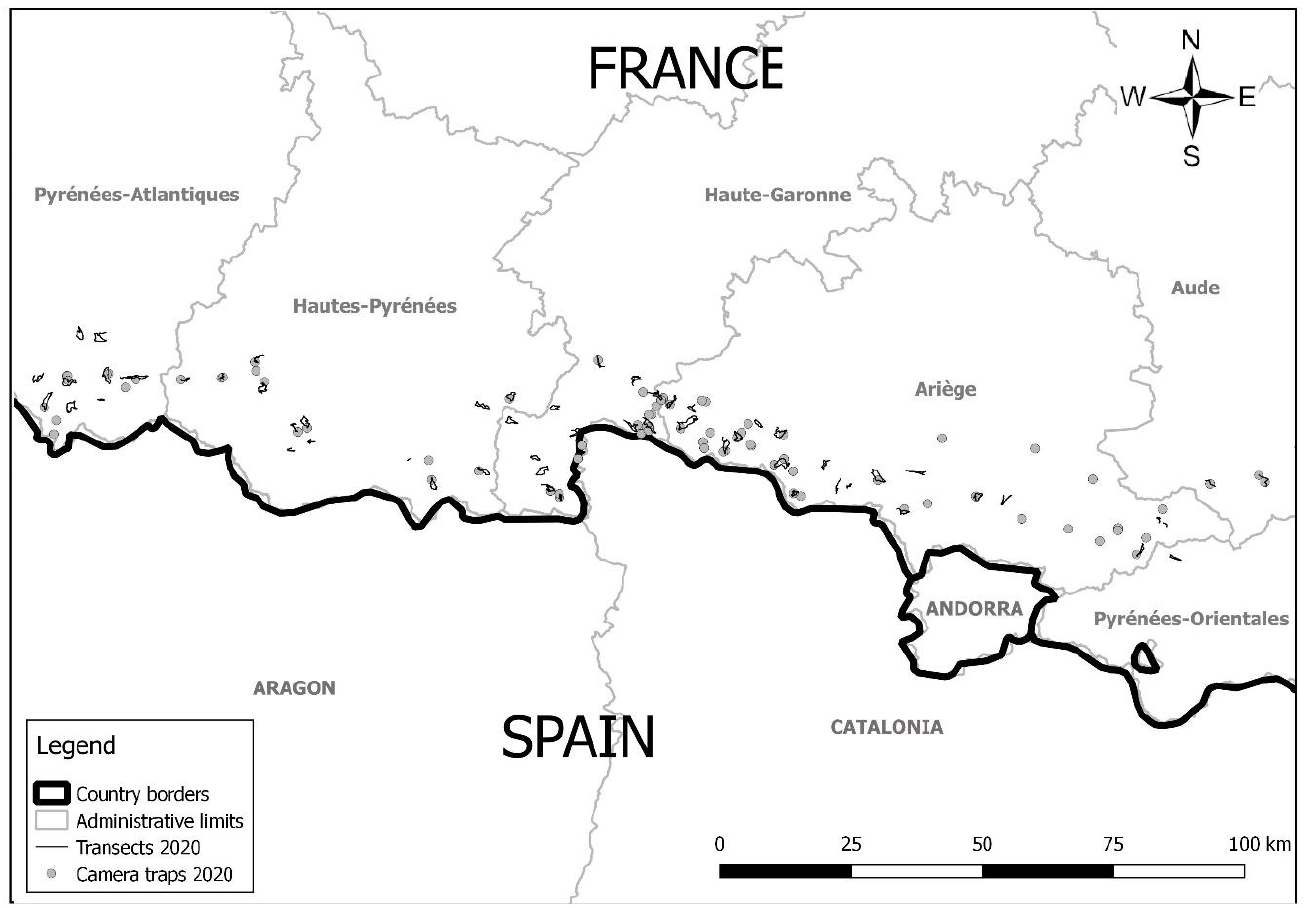
Map of the camera traps and transects used in 2020 in France within the framework of the systematic monitoring of the Pyrenean brown bear population.

