## Supplementary material for "Estimating abundance of a recovering transboundary brown bear population with capture-recapture models": Supp Material

**Supplementary Information of the paper entitled "Estimating abundance of a recovering transboundary brown bear population with capture-recapture models" by Vanpé et al.**

### Materials and Methods

*Genetic analyses from 2017 to 2020*

**Table S1.** Systematic monitoring effort in the French Pyrenees in terms of number of transects (including 6 hair traps per transect in average), total length of transects (km), number of camera traps, number of baited hair traps per year between 2008 and 2020.

| Year | # transects | total length of transects (km) | # camera traps | # baited hair traps |
|---|---|---|---|---|
| 2008 | 30 | 300* | 7 | 59 |
| 2009 | 36 | 360* | 10 | 73 |
| 2010 | 60 | 600* | 12 | 86 |
| 2011 | 68 | 615 | 26 | 90 |
| 2012 | 68 | 615 | 18 | 0 |
| 2013 | 48 | 426 | 49 | 0 |
| 2014 | 50 | 411 | 39 | 0 |
| 2015 | 44 | 358 | 40 | 0 |
| 2016 | 47 | 376 | 48 | 0 |
| 2017 | 53 | 414 | 45 | 0 |
| 2018 | 57 | 441 | 45 | 0 |
| 2019 | 56 | 424 | 59 | 0 |
| 2020 | 58 | 428 | 60 | 0 |

Note: * Estimated based on an average transect length of 10 km.

**Table S2.** Total number of validated non-invasive brown bear signs (e.g., scats, hair, tracks, visual observations, damages, photos / videos) collected in the Pyrenees, total number of validated brown bear samples (i.e. scats and hair) collected in the Pyrenees, number of samples (among collected sampled) genetically analysed by the French molecular laboratory LECA or ANTAGENE, number of brown bear samples (among analysed samples) successfully genotyped and number of different brown bear genotypes identified (among successfully genotyped samples) per year between 2008 and 2020.

| Year | # validated bear signs collected | # bear samples | # genetically analysed samples | # successfully genotyped samples | # different genotypes identified |
|---|---|---|---|---|---|
| 2008 | 743 | 210 | 125 | 73 | 11 |
| 2009 | 712 | 229 | 84 | 42 | 12 |
| 2010 | 939 | 323 | 167 | 106 | 15 |
| 2011 | 1152 | 518 | 209 | 122 | 15 |
| 2012 | 1239 | 521 | 224 | 153 | 15 |
| 2013 | 1318 | 521 | 137 | 77 | 14 |
| 2014 | 1243 | 571 | 193 | 96 | 21 |
| 2015 | 1567 | 870 | 152 | 110 | 24 |
| 2016 | 1854 | 874 | 179 | 137 | 32 |
| 2017 | 1394 | 569 | 134 | 105 | 34 |
| 2018 | 1625 | 601 | 158 | 109 | 35 |
| 2019 | 2450 | 830 | 314 | 209 | 41 |
| 2020 | 2783 | 1116 | 448 | 309 | 45 |
| TOTAL | 19019 | 7753 | 2524 | 1648 | 314 |

**Table S3.** Combination of microsatellite markers used in each PCR mix and type of fluorescent dye used for each microsatellite marker from 2017 to 2020.

| Mix | Locus name | Dye | Publication |
|---|---|---|---|
| A | UA03 | 6FAM | De Barba *et al*. 2017 |
| A | UA06 | 6FAM | De Barba *et al*. 2017 |
| A | UA25 | NED$^{TM}$ | De Barba *et al*. 2017 |
| A | UA67 | NED$^{TM}$ | De Barba *et al*. 2017 |
| A | UA64 | PET$^{TM}$ | De Barba *et al*. 2017 |
| A | UA63 | PET$^{TM}$ | De Barba *et al*. 2017 |
| A | UA16 | VIC$^{TM}$ | De Barba *et al*. 2017 |
| A | UA14 | VIC$^{TM}$ | De Barba *et al*. 2017 |
| B | UA17 | 6FAM | De Barba *et al*. 2017 |
| B | UA57 | 6FAM | De Barba *et al*. 2017 |
| B | UA51 | NED$^{TM}$ | De Barba *et al*. 2017 |
| B | UA65 | PET$^{TM}$ | De Barba *et al*. 2017 |
| B | UA68 | VIC$^{TM}$ | De Barba *et al*. 2017 |
| B | Our-ZFX | 6FAM | Bidon *et al*. 2013 |
| B | Our-318 | 6FAM | Bidon *et al*. 2013 |
| B | Our-SMCY | 6FAM | Bidon *et al*. 2013 |

**Table S4.** Summary statistics of the 58 different genotypes found in the Pyrenean brown bear population in 2020 for each of the 13 microsatellite loci provided by the allele frequency analysis of CERVUS software (Marshall et al. 1998).

| Locus | N | k | HObs | HExp | NE-I | NE-SI | PIC | F(Null) |
|---|---|---|---|---|---|---|---|---|
| UA03 | 58 | 4 | 0.655 | 0.622 | 0.215 | 0.495 | 0.549 | -0.0218 |
| UA06 | 58 | 4 | 0.724 | 0.657 | 0.173 | 0.467 | 0.6 | -0.0653 |
| UA14 | 58 | 4 | 0.759 | 0.705 | 0.144 | 0.437 | 0.645 | -0.0458 |
| UA16 | 58 | 6 | 0.414 | 0.461 | 0.328 | 0.604 | 0.424 | 0.0908 |
| UA17 | 58 | 3 | 0.517 | 0.497 | 0.308 | 0.581 | 0.442 | -0.0189 |
| UA25 | 58 | 5 | 0.483 | 0.427 | 0.364 | 0.629 | 0.392 | -0.0989 |
| UA51 | 58 | 4 | 0.603 | 0.537 | 0.269 | 0.551 | 0.483 | -0.0608 |
| UA57 | 58 | 3 | 0.552 | 0.45 | 0.399 | 0.627 | 0.354 | -0.1089 |
| UA63 | 57 | 6 | 0.719 | 0.694 | 0.146 | 0.442 | 0.639 | -0.0178 |
| UA64 | 58 | 2 | 0.534 | 0.492 | 0.381 | 0.601 | 0.369 | -0.0455 |
| UA65 | 58 | 4 | 0.621 | 0.595 | 0.246 | 0.516 | 0.513 | -0.0358 |
| UA67 | 58 | 3 | 0.517 | 0.571 | 0.266 | 0.533 | 0.488 | 0.0589 |
| UA68 | 58 | 5 | 0.724 | 0.734 | 0.121 | 0.417 | 0.68 | -0.0054 |
| MEAN | | 4.08 | 0.602 | 0.572 | 0.258 | 0.531 | 0.506 | -0.0289 |

Note: N: number of individuals typed, k: the number of alleles, Hobs: observed heterozygosity, Hexp: expected heterozygosity, NE-I: average exclusion probabilities for each locus for identity, NE-SI: average exclusion probabilities for each locus for sib identity, PIC: polymorphic information content, F(Null): the frequency of null alleles. The combined non-exclusion probabilities for identity and sib identity were $9.10^{-9}$ and 0.000235, respectively.

**Table S5.** Parameters of the model in which temporary emigration is random, survival is age-dependent and there is heterogeneity in the detection process, estimated using a Bayesian robust-design capture-recapture (CR) approach.

| | Mean | SD | Naive SE | Time-series SE |
|---|---|---|---|---|
| beta[1] | 8.52E-01 | 0.0402 | 0.8991 | 0.0014 |
| beta[2] | 9.33E-01 | 0.0333 | 0.0007 | 0.0009 |
| beta[3] | 9.46E-01 | 0.0175 | 0.0004 | 0.0004 |
| deviance | 2.31E+03 | 17.9349 | 0.4010 | 0.5163 |
| gamma | 6.62E-02 | 0.0218 | 0.0005 | 0.0006 |
| mean.p | 4.29E-01 | 0.0272 | 0.0006 | 0.0019 |
| pstar[1] | 8.50E-01 | 0.0188 | 0.0004 | 0.0009 |
| pstar[2] | 8.50E-01 | 0.0188 | 0.0004 | 0.0009 |
| pstar[3] | 8.50E-01 | 0.0188 | 0.0004 | 0.0009 |
| pstar[4] | 8.50E-01 | 0.0188 | 0.0004 | 0.0009 |
| pstar[5] | 8.50E-01 | 0.0188 | 0.0004 | 0.0009 |
| pstar[6] | 8.50E-01 | 0.0188 | 0.0004 | 0.0009 |
| pstar[7] | 8.50E-01 | 0.0188 | 0.0004 | 0.0009 |
| pstar[8] | 8.50E-01 | 0.0188 | 0.0004 | 0.0009 |
| pstar[9] | 8.50E-01 | 0.0188 | 0.0004 | 0.0009 |
| pstar[10] | 8.50E-01 | 0.0188 | 0.0004 | 0.0009 |
| pstar[11] | 8.50E-01 | 0.0188 | 0.0004 | 0.0009 |
| pstar[12] | 8.50E-01 | 0.0188 | 0.0004 | 0.0009 |
| sdeps | 2.10E-01 | 0.0214 | 0.0005 | 0.0007 |

Note: beta[i]: age-specific survival for age i (with 1: cubs, 2: subadults, 3: adults), gamma: probability of emigration; mean.p: mean detection probability, sdeps: SD of the random effect, pstar[j]: averaged detection over individuals for year j, with j ranging from 2008 to 2019.

**Table S6.** Comparison of the annual abundance of the Pyrenean brown bear population, estimated from Bayesian Pollock's robust design (PCRD) capture-recapture (CR) approach (with associated 95% Credible Interval), with Minimum Detected Size (MDS, total number of different individuals detected in the population during the year) and Minimum Retained Size (MRS, reassessment of the MDS in the light of the information collected in subsequent years) values from 2008 to 2020.

| Year | PCRD Estimate | 95% CI | MDS value | MRS value |
|---|---|---|---|---|
| 2008 | 13.0 | 12.8 - 13.3 | 16 | 15 |
| 2009 | 17.4 | 17.0 - 17.8 | 17 | 16 |
| 2010 | 16.3 | 15.9 - 16.7 | 18 | 20 |
| 2011 | 19.5 | 19.1 - 20.0 | 22 | 23 |
| 2012 | 23.9 | 23.4 - 24.4 | 22 | 24 |
| 2013 | 21.7 | 21.3 - 22.2 | 25 | 25 |
| 2014 | 26.0 | 25.5 - 26.7 | 31 | 31 |
| 2015 | 29.3 | 28.7 - 30.0 | 29 | 32 |
| 2016 | 41.2 | 40.4 - 42.2 | 39 | 41 |
| 2017 | 41.2 | 40.4 - 42.2 | 43 | 48 |
| 2018 | 39.1 | 38.3 - 40.0 | 40 | 52 |
| 2019 | 49.9 | 48.9 - 51.1 | 52 | 59 |
| 2020 | 66.2 | 64.8 - 67.8 | 64 | 68 |

Note: MRS count for 2020 is provisional and probably slightly underestimated.

**Table S7.** Evolution of the sex ratio of the Pyrenean brown bear population from 2008 to 2020 among all individuals and among adult only.

| | # adult males | # adult females | Adult sex ratio | # males | # females | Sex ratio |
|---|---|---|---|---|---|---|
| 2008 | 5 | 4 | 1.25 | 8 | 6 | 1.33 |
| 2009 | 6 | 5 | 1.20 | 8 | 8 | 1.00 |
| 2010 | 5 | 6 | 0.83 | 8 | 12 | 0.67 |
| 2011 | 6 | 6 | 1.00 | 8 | 14 | 0.57 |
| 2012 | 6 | 7 | 0.86 | 9 | 14 | 0.64 |
| 2013 | 6 | 9 | 0.67 | 9 | 14 | 0.64 |
| 2014 | 8 | 11 | 0.73 | 11 | 17 | 0.65 |
| 2015 | 7 | 12 | 0.58 | 13 | 19 | 0.68 |
| 2016 | 7 | 13 | 0.54 | 19 | 21 | 0.90 |
| 2017 | 8 | 14 | 0.57 | 21 | 26 | 0.81 |
| 2018 | 8 | 19 | 0.42 | 22 | 28 | 0.79 |
| 2019 | 11 | 21 | 0.52 | 23 | 28 | 0.82 |
| 2020 | 12 | 23 | 0.52 | 29 | 29 | 1.00 |

**Figure. S1.** Map of the camera traps and transects used in 2020 in France within the framework of the systematic monitoring of the Pyrenean brown bear population.

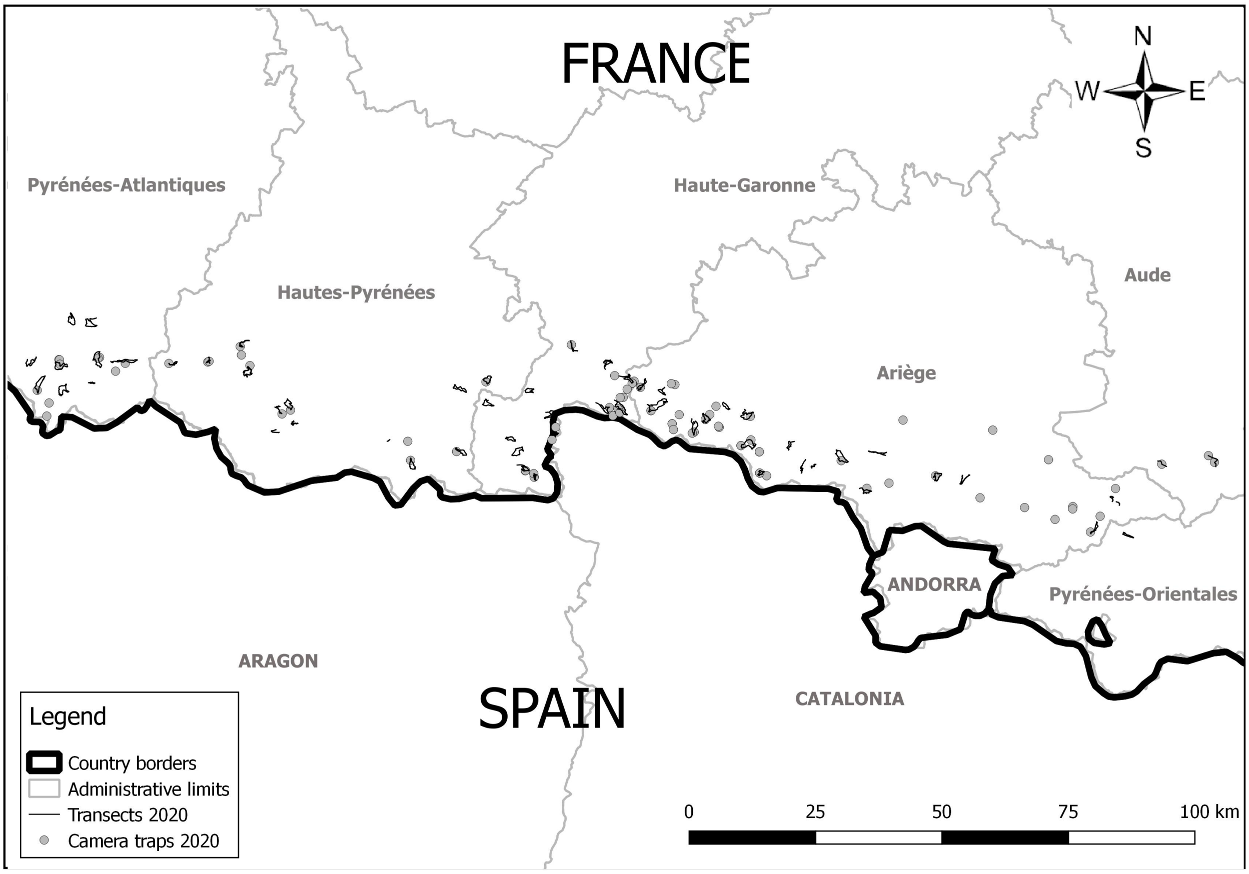
